# Graph Properties of the Adult *Drosophila* Central Brain

**DOI:** 10.1101/2020.05.18.102061

**Authors:** Louis K. Scheffer

**Author notes:** Corresponding author: Lou Scheffer.

## Abstract

The recent *Drosophila* central brain connectome offers the possibility of analyzing the graph properties of the fly brain. Crucially, this connectome is dense, meaning all nodes and links are represented, within the limits of experimental error. We consider the connectome as a directed graph with weighted edges. This enables us to look at a number of graph properties, compare them to human designed logic systems, and speculate on how this may affect function. We look at input and output distributions, randomness of wiring, differences between compartments, path lengths, proximity of strong connections, known computational structures, electrical response as a function of compartment structure, and evidence for efficient packing.

## 1 Introduction

The connectome we analyze[1] is a dense reconstruction of a portion of the central brain (referred to here as the hemibrain) of the fruit fly, *Drosophila melanogaster*, as shown in Figure 1. This connectome contains around 25,000 neurons with about 20 · 10^6^ chemical synapses between them, plus portions of many other neurons truncated by the boundary of the data set (details in Figure 1 below).

**Figure 1:**
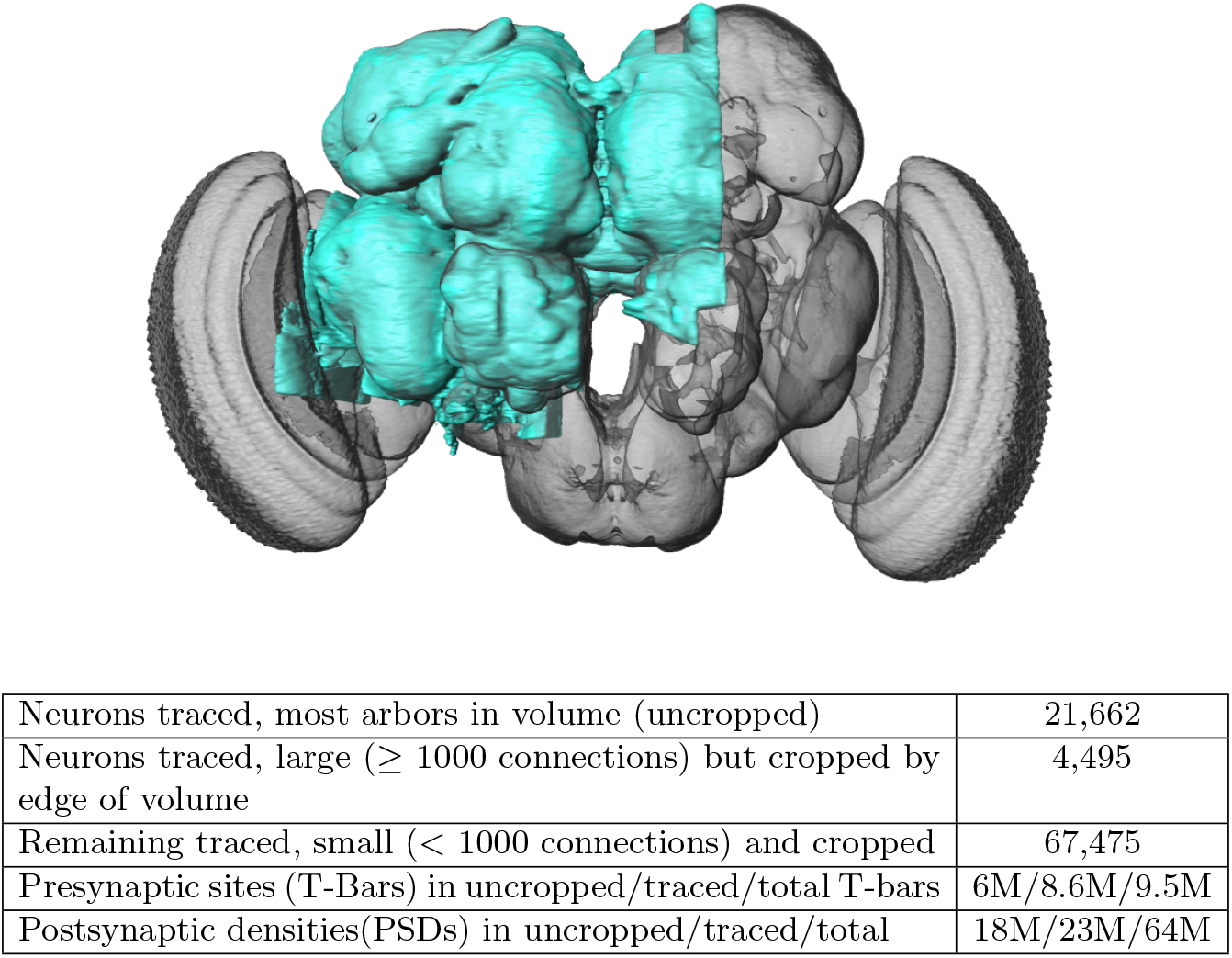
The hemibrain and some basic statistics. The highlighted area shows the portion of the central brain that was imaged and reconstructed, superimposed on a grayscale representation of the entire *Drosophila* brain. For the table, a neuron is traced if all its main branches within the volume are reconstructed. A neuron is considered uncropped if most arbors (though perhaps not the soma) are contained in the volume. Others are considered cropped. Note: 1) our definition of cropped is somewhat subjective; 2) the usefulness of a cropped neuron depends on the application; and 3) some small fragments are known to be distinct neurons. For simplicity, we will often state that the hemibrain contains ≈25K neurons.

Here we consider the fly brain as a directed graph where neurons are nodes and synaptic connections are edges. The edges have weights equal to the number of synapses connecting the nodes in the specified direction. We equate this weight with the strength of a connection, though we know this to be an approximation as many other factors (such as the transmitter and receptors used, and the underlying biochemistry) also affect the biological strength. We generate graph metrics for each computational compartment, and for the sample as a whole. Some of these analyses have already been covered in the initial connectome paper[2]; others are new here. They are combined here to put the related analyses in one place.

The graph metrics we consider are:

- N, the number of nodes.
- L, the number of links.
- 〈*n*〉, the average number of links per node (*L*/*N*).
- p, the probability that a link exists, = *L*/(*N* * (*N* + 1)).
- D, the diameter, or longest of the shortest paths between any two nodes in the graph.
- |*CC*|, the size of connected components. This is computed on the undirected graph as a directed graph gives ambiguous results.
- 〈*s*〉, the average connection strength, with three distinctions - overall, reciprocal connections, and non-reciprocal connections.

Since there is no consensus on the importance of small connections, all these may be measured a function of T, the threshold, which is the lowest weight connection used in the computation.

## 2 Analysis

There are many different ways we can analyze the brain graph.

### 2.1 Overall statistics

Over our entire sample, we have 22000 traced neurons with about 3.4M connections between them. It is, however, a subset of the whole brain and hence the overall brain statistics we note here are preliminary. We observe a wide range of upstream and downstream partners. The lowest number of partners is zero, almost surely a result of sample truncation. The largest numbers are 5044 inputs and 2708 outputs. These numbers are almost surely underestimates, since the largest neurons typically span the entire brain and our sample comprises only a portion. In our sample, each neuron averages 153 downstream and 153 upstream partners (over the whole sample, these two numbers must be the same).

The fly brain contains one connected component containing at least 99.9% of all neurons, and has a diameter of 12, so the path from any one of these neurons to any other passes through at most 12 synapses.

### 2.2 The fly brain is not wired randomly

The brain is not wired randomly. This seems obvious, and has been widely reported in previous studies, but is confirmed by the graph of degree distribution. Here we compare the observed degree distribution compared to what we would expect from a random graph with the same number of nodes and the same number of links per node. These appear very different, as shown in Figure 2, and a Kolmogorov-Smirnov test confirms this, with an extremely low chance they are drawn from the same distribution (*p* < 10^−300^).

**Figure 2:**
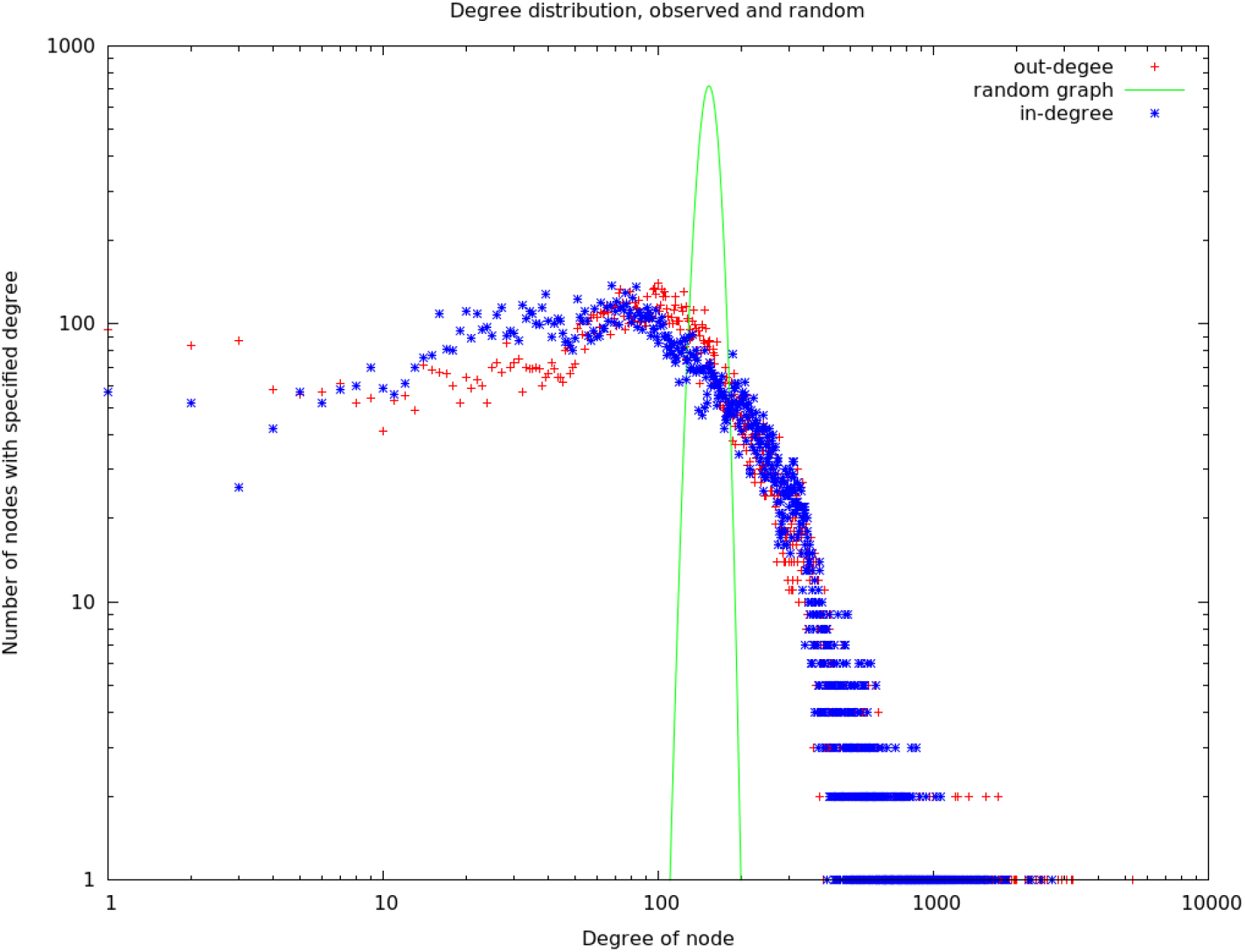
The observed degree distribution. The random graph is the degree distribution expected for the same number of nodes and links, but where the links are chosen at random from all possible links.

### 2.3 Compartment statistics

One analysis enabled by a dense whole-brain reconstruction involves the comparison between the circuit architectures of different brain areas within a single individual.

The compartments vary considerably. Table 1 shows the connectivity statistics of compartments that are completely contained within the volume, have at least 100 neurons, and have the largest or smallest value of various statistics. Across regions, the number of neurons varies by a factor of 74, the average number of partners of each neuron by a factor of 36, the network diameter by a factor of 4, the average strength of connection between partner neurons by a factor of 5, and the fraction of reciprocal connections by a factor of 5. The average graph distance between neurons is more conserved, differing by a factor of only 2.

**Table 1:** Regions with minimum or maximum characteristics, picked from those regions lying wholely within the reconstructed volume and containing at least 100 neurons. Yellow indicates a minimum value; green a maximal value. N is the number of neurons in the region, L the number of connections between those neurons, 〈*k*〉 the average number of partners (in the region), D the network diameter, 〈str〉 the average connection strength, broken up into non-reciprocal and reciprocal. fracR is the fraction of connections that are reciprocal, and AvgDist is the average number of hops (one hop corresponding to a direct synaptic connection) between any two neurons in the compartment.

| Name | N | L | $\langle k \rangle$ | D | $\langle \text{str} \rangle$ | $\langle \text{non-r} \rangle$ | $\langle r \rangle$ | fracR | AvgDist |
| --- | --- | --- | --- | --- | --- | --- | --- | --- | --- |
| MB(R) | 3430 | 573664 | 167.249 | 8 | 3.270 | 3.076 | 3.383 | 0.632 | 2.189 |
| bL(R) | 1159 | 108134 | 93.299 | 8 | 2.019 | 1.855 | 2.122 | 0.613 | 2.090 |
| EB | 518 | 58793 | 113.500 | 4 | 10.053 | 4.627 | 12.172 | 0.719 | 1.756 |
| PLP(R) | 6689 | 224985 | 33.635 | 16 | 2.692 | 2.403 | 3.685 | 0.226 | 3.170 |
| SNP(R) | 9121 | 775474 | 85.021 | 13 | 2.987 | 2.515 | 4.492 | 0.239 | 2.748 |
| RUB(L) | 123 | 576 | 4.683 | 6 | 7.682 | 2.852 | 20.686 | 0.271 | 2.743 |
| EPA(R) | 1468 | 18199 | 12.397 | 13 | 2.171 | 2.098 | 2.644 | 0.134 | 3.496 |

### 2.4 Paths in the fly brain are short

Neurons in the fly brain are tightly interconnected, as shown in Figure 3, which plots what fraction of neuron pairs are connected as a function of the number of interneurons between them. Three quarters of all possible pairs are connected by a path with less than three interneurons, even when only connections with ≥ 5 synapses are included. If weaker connections are allowed, the paths become shorter yet. These short paths and tight coupling are very different from human designed systems, which have much longer path lengths connecting node pairs. As an example, a standard electrical engineering benchmark (S38584 from [3]) is shown alongside the hemibrain data in Figure 3A-B. The connection graph for this example has roughly the same number of nodes as the graph of the fly brain, but pair-to-pair connections involve paths more than an order of magnitude longer – a typical node pair is separated by 60 intervening nodes. This is because a typical computational element in a human designed circuit (a gate) connects only to a few other elements, whereas a typical neuron reeives input from, and sends outputs to, hundreds of other neurons.

**Figure 3:**
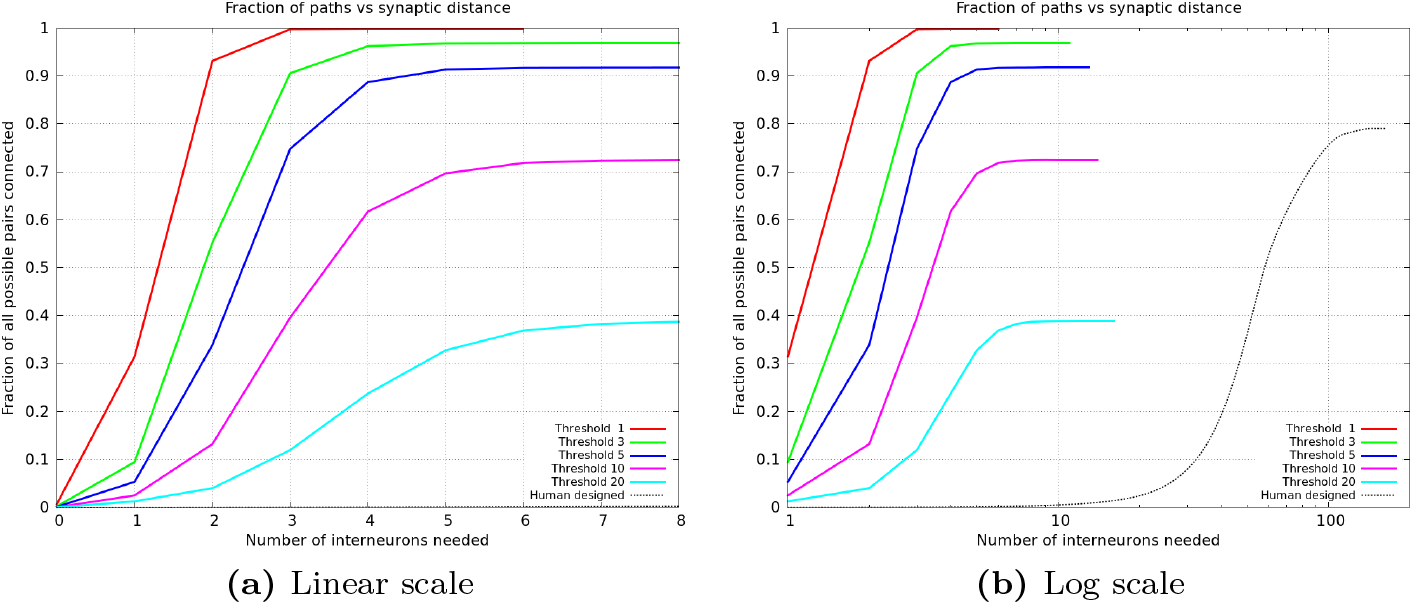
Plots of the percentage of pairs connected (of all possible) versus the number of interneurons required. (a) shows the data from the whole hemibrain, for up to 8 interneurons. (b) is a much wider view of the same data, shown on a log scale so the curve from a human designed system is visible.

### 2.5 No single input contributes strongly

No single input dominates the input to a neuron. As shown in Figure 4, for the median neuron, the strongest input contributes only about 4% of the total incoming synaptic strength. For almost all neurons, > 90%, the strongest input contributes less than 10% of all input. Very nearly the same distribution holds for outputs - each neuron typically drives many others, with none dominating in strength. This distribution is in contrast to that of man-made systems, where fan-in of 2-10 is considered typical[4][5], and a fan-in of 64 is considered quite high[6]. In contrast typical neuron in our sample has an average fan-in of 153.

**Figure 4:**
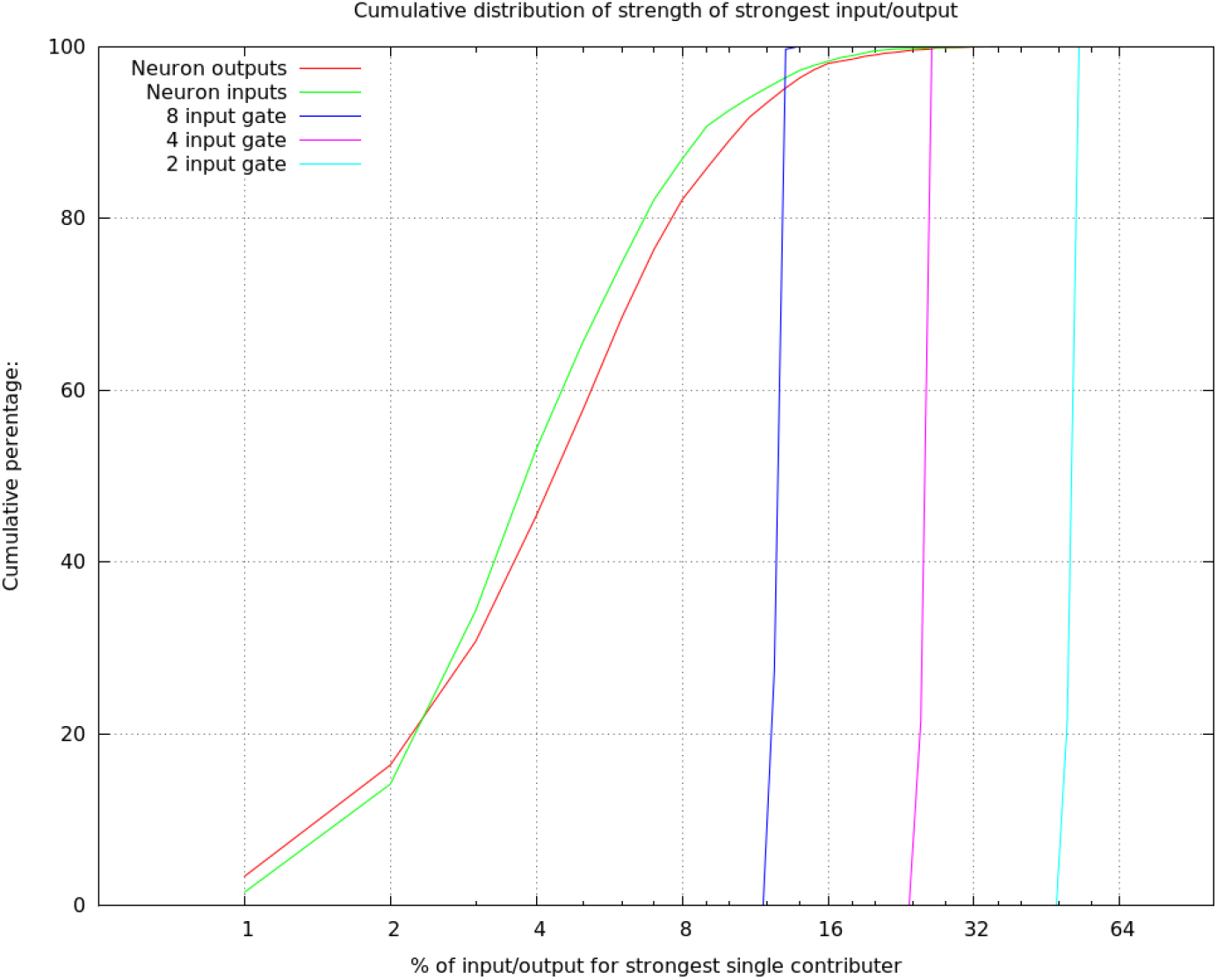
Plot of the percentage of input contributed by the strongest single input. This is cumulative over all neurons in the data set. Also shown, for comparison, are three cases from human designed systems - a logic gate with 2, 4, or 8 inputs. The graphs of human systems are more correctly step functions, but drawn slightly tilted for visibility.

### 2.6 Distribution of connection strength

The distribution of connection strengths has been studied in mammalian tissue, looking at specific cell types in specific brain areas. These findings, such as the log-normal distribution of connection strengths in rat cortex, do not appear to generalize to flies. Assuming the strength of a connection is proportional to the number of synapses in parallel, we can plot the distribution of connection strengths, summing over the whole central brain, as shown in Figure 5. We find a nearly pure power law with an exponential cutoff, very different from the log-normal distribution of strengths found by Song[7] in pyramidal cells in the rat cortex, or the bimodal distribution found for pyramidal cells in the mouse by Dorkenwald[8]. However, we caution that these analyses are not strictly comparable. Even aside from the very different species examined, the three analyses differ. Both Song and Dorkenwald looked at only one cell type, with excitatory connections only, but one looked at electrical strength while the other looked at synapse area as a proxy for strength. In our analysis, we use synapse count as a proxy for connection strength, and look at all cell types, including both excitatory and inhibitory synapses.

**Figure 5:**
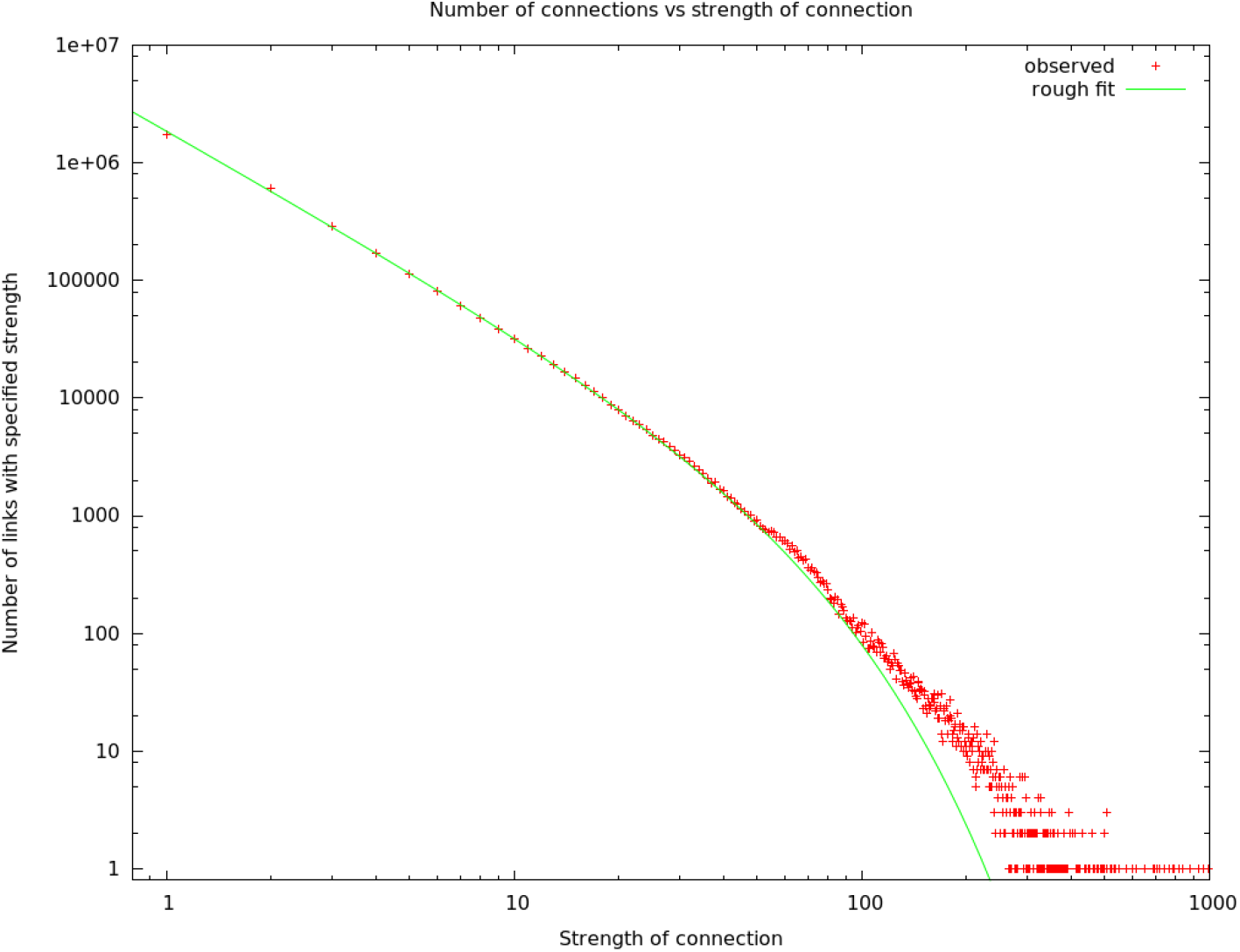
The number of connections with a given strength. Up to a strength of 100, this is well described by a power law (exponent −1.67) with exponential cutoff (at N=42).

### 2.7 Strong connections are not uniformly distributed

Song[7] observed the general trend that strong edges cluster together in a subset of mammalian cells. We find the same trend holds more broadly, and can quantify it using connected components, as shown in Table 2 and Figure 6..

**Figure 6:**
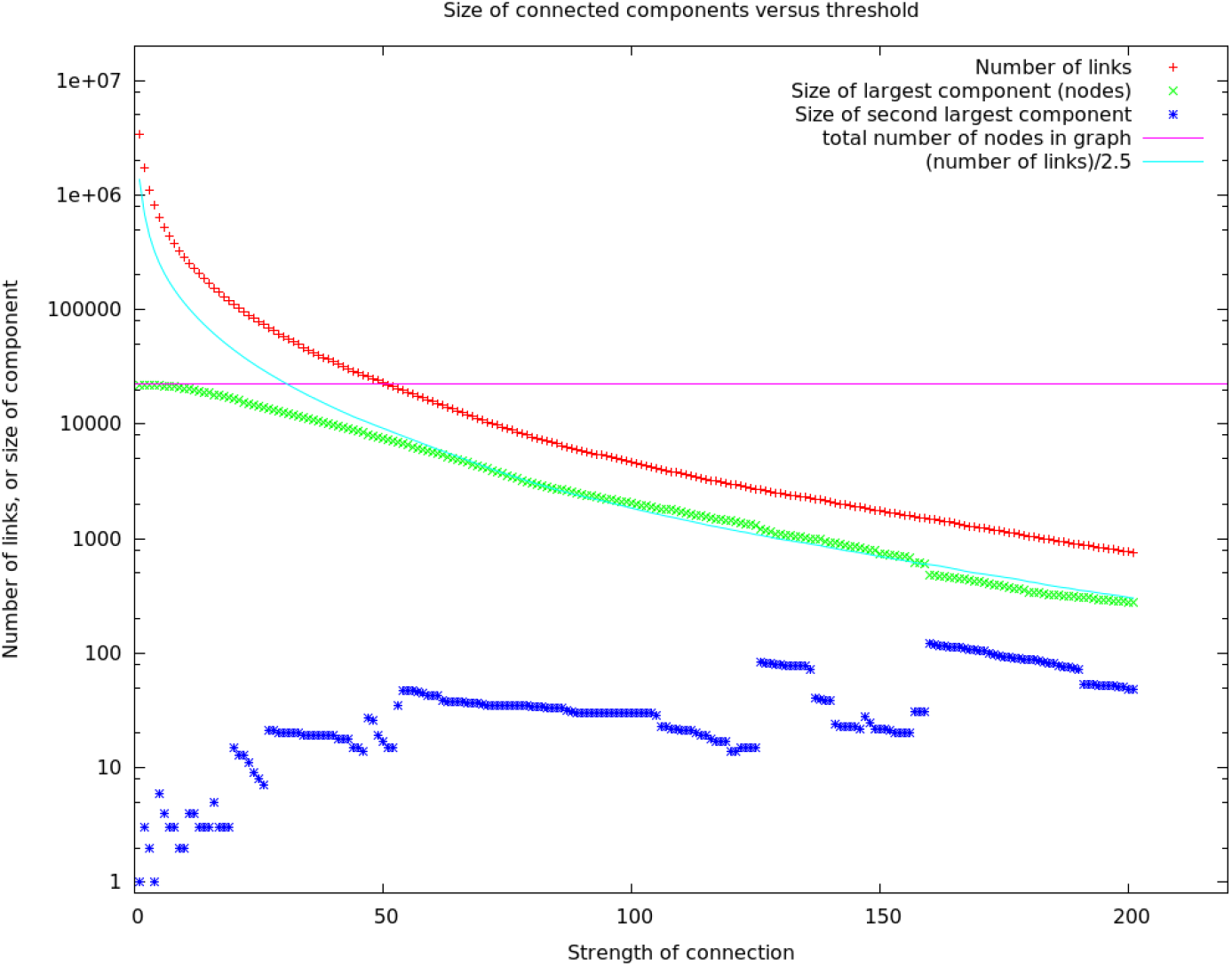
Size of connected components versus threshold.

**Table 2:** Strong connections occur in small groups

| Threshold | Number of edges ≥ threshold | Size of connected component | % of edges in selected component |
| --- | --- | --- | --- |
| 1 | 3411204 | 22178 | >99.9% |
| 51 | 22159 | 7360 | 98.8% |
| 101 | 4509 | 1991 | 92.3% |
| 151 | 1721 | 736 | 76.3% |
| 201 | 751 | 276 | 64.2% |

By increasing the threshold T, we can reduce the total number of edges until there are fewer edges than nodes (which happens at T = 51 and higher). If the strong edges were randomly distributed among the graph, at this point it would be expected to transition to a state where there are no large connected components[9]. But this is not what we observe - even at T = 101, where only 4500 edges remain, there is a connected component of 1900 nodes (<10% of the nodes) that contains more than 90% of the edges. At all strength thresholds, the connected components are dominated by one large component.

One feature we do not see is a division into two large connected components. This could potentially happen, for example, if each side of the brain formed a component, and the two sides were lightly linked. One possible explanation is that the two sides of the brain are not equally represented in our sample, as shown in Figure 1. However, we think a more likely explanation is that the cross brain connections are among the strongest links. These hypotheses can be tested once we have a full brain connectome, work that is in progress now.

The large component is almost as large as it can be. It cannot exceed the number of nodes, nor the number of links (plus one). The size of the largest component, for thresholds > 50, is observed to be roughly equal to the number of links divided by 2.5. Since the largest component dominates, this implies each node in this component has about 2-2.5 links, on the average, independent of threshold.

### 2.8 Small Motifs

As mentioned earlier, there have been many studies of small motifs, usually involving limited circuits, cell types, and brain regions. We emphatically confirm some traditional findings, such as the over-representation of reciprocal connections. We observe this in all brain regions and among all cell types, confirming similar findings in the antennal lobe[10]. This can now be assumed to be a general feature of the fly’s brain, and possibly all brains. In the fly, the incidence varies somewhat by compartment, however, as shown in Table 1.

### 2.9 Large motifs

We define a large motif as a graph structure that involves every cell of an abundant type (N ≥ 20). The most tightly bound motif is a clique, in which every cell of a given type is connected to every other cell of that type, with synapses in both directions. Such connections, as ilustrated in Figure 7(a), are extremely unlikely in a random wiring model. Consider, for example, the clique of R4d_b cells found in the ellipsoid body, as shown in Table 3. In the ellipsoid body, two cells are connected with an average probability of 0.19. Therefore the odds of finding all 600 possible connections between R4d_b cells, assuming a random wiring model, is 0.19^600^ ≈ 10^-432^.

**Figure 7:**
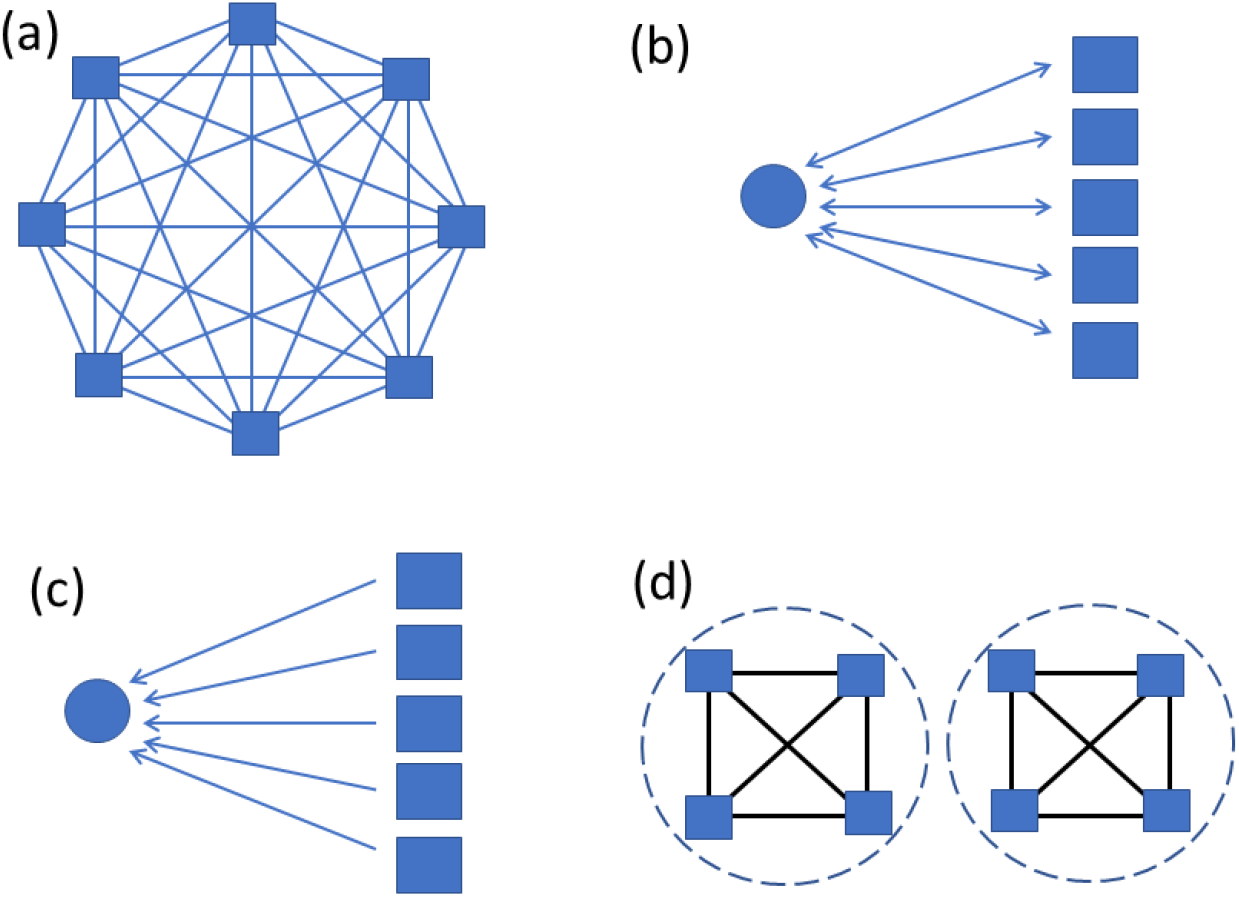
Large motifs searched for. Squares represent abundant types with at least 20 instances. Circles represent sparse types with at most two instances. Panel (a) shows a clique, where all possible connections are present. (b) shows bidirectional connections between a sparse type and all instances of an abundant type. (c) show unidiectional connections from all of an abundant type to a sparse type. (d) illustrates a cell type that does not form a clique overall, but does within each of two compartments.

**Table 3:** Cliques and near-cliques in the hemibrain data. To be included, a cell type must have at least 20 cell instances, 90% or more of which connect both to and from at least 90% of all cells of the same type. Coverage is the fraction of all possible edges in the clique that are present. Average strength is the average number of synapses in each connection. Synapses is the total number of synapses in the clique.

| Type | Region | Cells | Coverage | Avg. Strength | Synapses |
| --- | --- | --- | --- | --- | --- |
| KCab-p | MB | 55/62 | 3552/3782 | 5.04 | 17899 |
| Delta7_a | PB, CX | 32/32 | 992/992 | 13.79 | 13683 |
| R4d_b | EB, CX | 25/25 | 600/600 | 54.94 | 32961 |
| R5 | EB, CX | 20/20 | 380/380 | 26.62 | 10114 |
| R3m | EB, CX | 22/22 | 462/462 | 24.32 | 11238 |
| R3d_a | EB, CX | 20/20 | 377/380 | 28.46 | 10729 |
| PFNa | NO(R) | 29/29 | 811/812 | 6.73 | 5459 |
| PFNa | NO(L) | 29/29 | 811/812 | 7.22 | 5858 |
| PFNd | NO(R) | 20/20 | 377/380 | 7.67 | 2891 |
| PFNd | NO(L) | 20/20 | 378/380 | 7.59 | 2869 |

In the fly’s brain, large cliques occur in only a few cases, as shown in Table 3. All true cliques are in the central complex, with a near-clique among the KCab-p cells of the mushroom body. The cells of type PFNa form an interesting case. There are 58 such cells, 29 on each side. They do not form a clique as shown in Figure 7(a), as there are few connections between the opposite sides. But within each side, the 29 cells on that aside form a clique, as shown in Figure 7(d).

Presumably connections within the clique must be inhibitory, since with such strong connections and numerous short loops, positive feedback would soon lead to runaway excitation and saturation. A natural speculation is these cells implement a “winner takes all” circuit, with at most one cell in each clique active and inhibiting all others. This structure seems expensive in terms of the number of synapses, and their energy demands, so it presumably supports a function important to the fly’s survival that cannot be easily undertaken by a more parsimonious structure.

The next most tightly bound motifs are individual cells that connect both to and from all cells of a given type, but are themselves of a different type. This is illustrated in Figure 7(b). Such a motif is often speculated to be a gain or sparseness controlling circuit, where the single neuron reads the collective activation of a population and then controls their collective behavior. A well known example is the APL neuron in the mushroom body, which connects both to and from all the Kenyon cells, and is thought to regulate the sparseness of the Kenyon cell activation.

We search for this motif by looking at cells with few instances (one or two) connecting bidirectionally to almost all cells (at least 90%) of an abundant type (N >= 20). We find this motif in three regions of the brain – it is common in the CX (73 different cells overseeing 22 cell types), the optic lobe circuits (19 cells overseeing 14 types), and somewhat in the MB (12 types overseeing 9 types). Spreadsheets containing these cell types, who they connect to, and the numbers and strengths of their connections are found in the supplementary data. We only analyze the optical circuits here, since the mushroom body and central complex are the subjects of companion papers. We observe three variations on this motif - a single cell conected to all of a type (Figure 8(a), found 5 times), a single cell with bi-directional connections to many types (Figure 8(b), found once), and multiple cells all connected bidirectionally to a single type (Figure 8(c), found 3 times. We find one circuit that is a combination: There is one cell that connects bidirectionally to all the LC17 neurons, and then a higher order cell that connects bidirectionally to a larger set (LPLC1, LPLC2, LLP1, LPC1, and LC17). In this case these are all looming-sensitive cells and hence these circuits may regulate the features of the overall looming responses. It is tempting to speculate that the more complex structures of Figure 8 (b) and (c) arose from the simpler structures of (a) through cell type duplication followed by divergence, but the connectomes of many more related species will be needed before this argument could be made quantitative.

**Figure 8:**
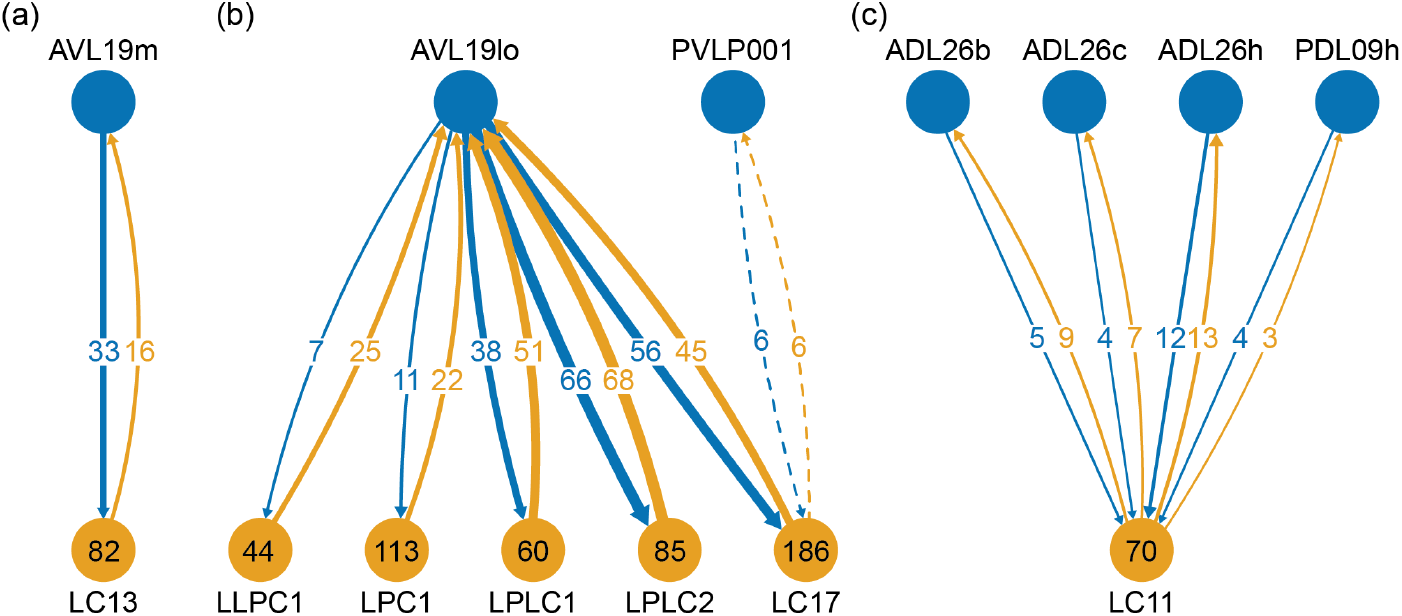
One to many motifs found in the optic circuits. Individual neurons and their names are shown at the top of the diagram. Cell type names, at the bottom of the diagram, represent cells with many instances, with the number of instances shown inside. The arrows show the average synapse count of each connection type. (a) shows an example of the most common case. Here one cell, AVL19m, has bidirectional connections to all cells of type LC13. (b) shows a single cell with exhaustive connections to several types. (c) shows an alternative motif where several cells form these one-to-many connections. For clarity the cell names have been truncated, with the suffix _pct (for putative cell type) removed.

The least tightly bound large motif is a cell that connects either to or from (but not both) all cells of a given type, as shown in Figure 7(c). Examples include the mushroom body output neurons[11]. This is a very common motif, found in many regions. We find more than 500 examples of this in the fly’s brain.

### 2.10 Brain regions and electrical response

How does the compartmentalization of the fly brain affect neural computation? In a few cases this has been established. For example, the CT1 neuron performs largely independent computations in each branch[12], whereas estimates show that within the medulla, the delays within each neuron are likely not significant for single column optic lobe neurons, and hence the neurons likely perform only a single computation[13].

Our detailed skeleton models allow us to construct electrical models of neurons. In particular, to look more generally at the issues of intra– vs inter– compartment delays and amplitudes, we can construct a linear passive model for each neuron. Our method is similar to that elsewhere[14], except that instead of using right cylinders, we represent each segment of the skeleton as a truncated cone. This is then used to derive the axonic resistance, the membrane resistance, and membrane capacitance for each segment. To analyze the effect of compartment structure on neuron operation, we inject the neuron at a postsynaptic density (input) with a signal corresponding to a typical synaptic input (1 nS conductance, 1 ms width, 0.1 ms rise time constant, 1 ms fall time constant, 60 mV reversal potential). We then compute the response at each of the T-bar sites (outputs). Since the synapses, both input and output, are annotated by the brain region that contains them, this allows us to calculate the amplitudes and delays from each synapse (or a sample of synapses) in each compartment to each output synapse in all other compartments.

In general, we find the ROI structure of the neuron is clearly reflected in the electrical response. Consider, for example, the EPG neuron (Figure 9(a)) with arbors in the ellipsoid body, the protocerebral bridge, and the gall. Figure 10(a) shows the responses to synaptic input in the gall. Within the gall, the delays are very short, and the amplitude relatively high and variable, depending somewhat on the input and output synapse within the gall. From the gall to other regions the delays are longer (typically a few milliseconds) and the amplitudes much smaller and nearly constant, largely independent of the exact transmitting and receiving synapse. There is a very clean separation between the within-ROI and across-ROI delays and amplitudes, as shown in Figure 10(a). The same overall behavior is true for inputs into the other regions - short delays and strong responses within the ROI, with longer delays and smaller amplitudes to other compartments.

**Figure 9:**
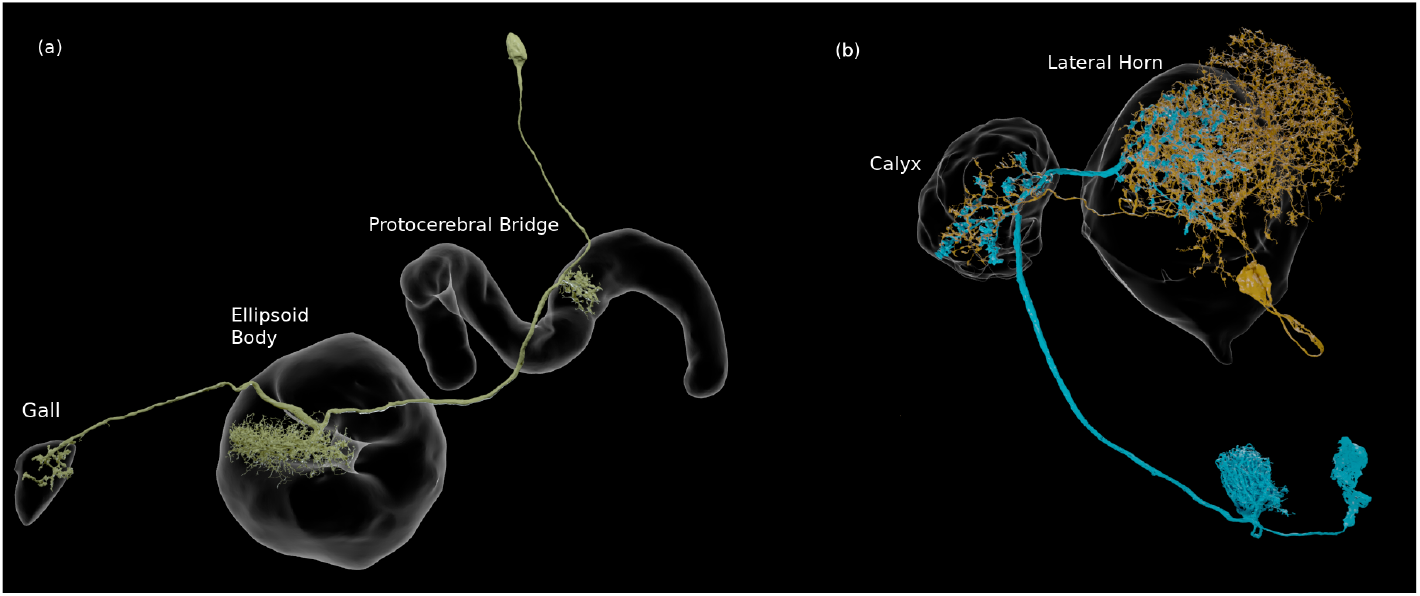
(a) An EPG neuron, with arbors in three compartments. (b) Two neurons that connect in more than one ROI, in this case the calyx and the lateral horn. They are each pre- and postsynaptic to each other in both compartments.

**Figure 10:**
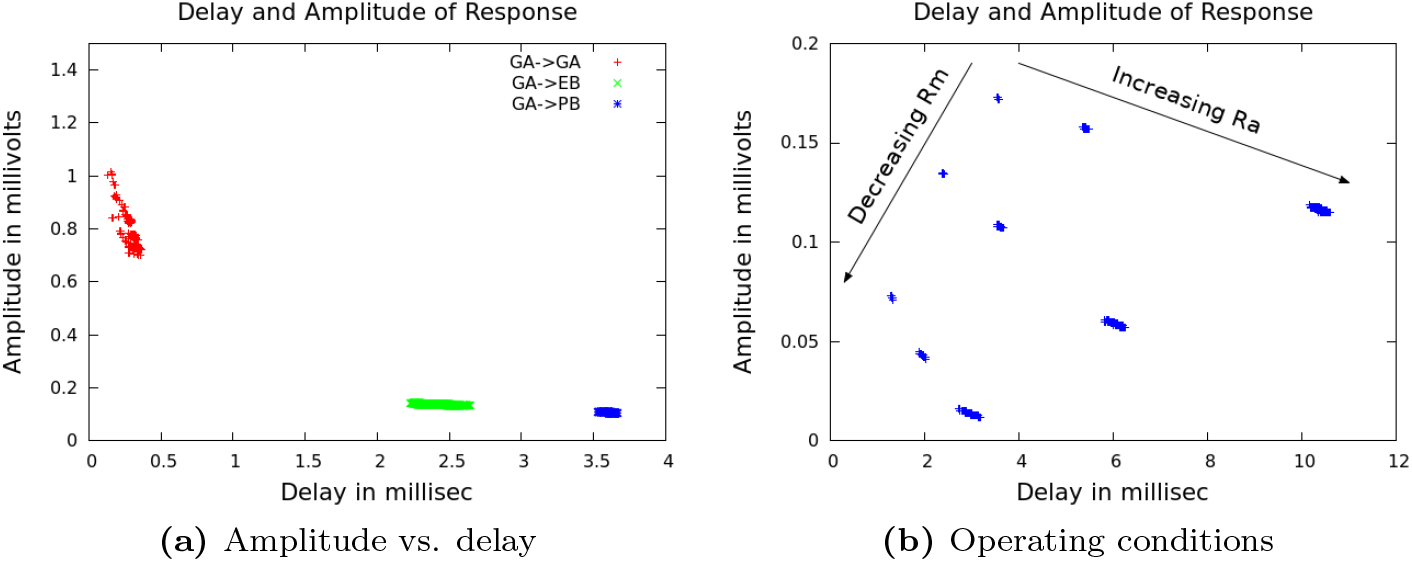
(a) The linear response to inputs in the gall(GA) for an EPG neuron, which also has arbors in the ellipsoid body(EB) and the protocerebral bridge (PB). Each point in the modeled plot shows the time each response reached its peak amplitude (the delay), and the amplitude at that time, for an input injected at one of the PSDs in the Gall. (b) Delays and amplitudes for gall to PB response, for all combinations of three values of cytoplasmic resistance *R_A_* and three values of membrane resistance *R_M_*.

This simple pattern motivates a model that describes delays and amplitudes not as a single number, but as NxN matrix, where N is the number of ROIs. Each row contains the estimated amplitude and delay, measured in each compartment, for a synaptic input in the given compartment. This gives a much improved estimate of the linear response. For the example EPG neuron above, with nominal values for *R_a_, R_m_*, and *C_m_*, if we represent all delays by a single number then the standard deviation of the error is 0.446 ms. If instead we represent the delays as a 3×3 matrix indexed by the compartment, the average error is 0.045 ms, for 10x greater accuracy. Similarly, the average error in amplitude drops from 0.168 mv to 0.021 mv, an eightfold improvement. While the improvement in error will depend on the neuron topology, in all cases it will be more accurate than a point model, for relatively little increase in complexity.

The absolute values of delay and amplitude are strongly dependent on the electrical parameters of the cell, however. A wide range of electrical properties have been reported in the fly literature (see Table 4) and it is plausible that these vary on a cell-to-cell basis. In addition, gap junctions, which are not included, may affect the value of *R_m_*. In light of this variation, we simulate with minimum, medium, and maximal values of *R_a_* and *R_m_*, for a total of 9 cases, as shown in Figure 10(b). All are needed since the resistance parameters interact non-linearly. We fix the value of *C_M_*, at 0.01 F/m^2^ since this value is determined by the membrane thickness and is not expected to vary from cell to cell[15]. The results over the parameter range are shown in Figure 10(b) for the case of the EPG neuron above for delay from the gall to the PB. The intra-ROI and between-ROI values are well separated for any value of the parameters (not shown).

**Table 4:** Values reported in the literature

| Reference | $R_a, \Omega \cdot \text{m}$ | $R_m, \Omega/\text{m}^2$ | $C_m, \text{F}/\text{m}^2$ |
| --- | --- | --- | --- |
| Borst[16], CH cells | 0.60 | 0.25 | 0.015 |
| Borst[16], HS cells | 0.40 | 0.20 | 0.009 |
| Borst[16], VS cells | 0.40 | 0.20 | 0.008 |
| Gouers[17], DM1 cell 1 | 1.62 | 0.83 | 0.026 |
| Gouers[17], DM1 cell 2 | 1.02 | 2.04 | 0.015 |
| Gouers[17], DM1 cell 3 | 2.66 | 2.08 | 0.008 |
| Gouers[17], dendrite 1 | 2.44 | 1.92 | 0.008 |
| Gouers[17], dendrite 2 | 2.66 | 2.08 | 0.008 |
| Gouers[17], dendrite 3 | 3.11 | 2.64 | 0.006 |
| Cuntz[18], HS cells | 4.00 | 0.82 | 0.006 |
| Meier[12], CT1 cells | 4.00 | 0.80 | 0.006 |

Programs that deduce synaptic strength and sign by fitting a computed response to a connectome and measured electrical or calcium imagindg data[19] may at some point require estimates of the delays within cells. If this is required, the above results suggest this could be accomplished with reasonable accuracy with a ROI-to-ROI delay table and 2 additional parameters per neuron, *R_A_* and *R_M_*. This is relatively few new parameters in addition to the many synaptic strengths already fitted.

A number of neurons have parallel connections in separate ROIs (see Figure 9(b)). This motif is common in the fly’s brain – about 5% of all connections having a strength ≥ 6 are spread across two or more non-adjacent ROIs. Given the increased delays and lower amplitudes of cross-compartment responses, this type of interaction differs electrically from those in which all connections are contained in a single ROI. A point neuron model cannot generate an accurate response for such connections – a synapse in region A will result in a fast response in A and a slower, smaller response in B, and vice versa, even though both of these events involve communication between the same two neurons. It is not known if this configuration has a significant influence on the neuron’s operation.

From these models we conclude (a) the compartment structure of the fly brain shows up directly in the electrical response of the neurons. (b) the compartment structure, though defined anatomically, matches that of the electrical response. From the clear separation in Figure 10, it is likely that the same compartment definitions could be found starting with the electrical response, though we have not tried this. (c) These results suggest a low dimensional model for neural operation, at least in the linear region. A small region-to-region matrix can represent the delays and amplitudes well. (d) Absolute delays depend strongly (but in a very predicable manner) on the values of axial and membrane resistance, which can vary both from animal to animal and from cell to cell. (e) Neurons that have a parallel connections in separate ROIs have a different electrical response than they would have with the same total number of synapses in a single ROI.

### 2.11 Rent’s rule analysis

Rent’s rule[20] is an empirical observation that in human designed computing systems, when the system is packed as tightly as possible, at every level of the hierarchy the required communication (the number of pins) scales as a power law of the amount of contained computation, measured in gates. Rent’s rule is an observed relationship, not derived from underlying theory, and the relationship is not exact and still contains scatter. A biological equivalent might be the observation that brain size tends to vary as a power law of body size[21], across a wide range of species occupying very different ecological and behavioral niches. Rent’s rule is roughly true over many orders of magnitude in scale, and for almost every system in which it has been measured. Somewhat surprisingly, Rent’s rule applies almost independently of the function performed by the computation being performed, and at every level of a hierarchical system. It also applies whether the compactness criterion is minimization of communication (partitioning) or physical close packing.

Rent’s rule is expressed as

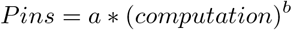

where *a* is a scale factor (typically in the range 1-4), and *b* is the ‘Rent exponent’ describing how the number of connections to the compartment varies as a function of the amount of computation performed in the compartment. The Rent exponent has a theoretical range of 0.0 to 1.0, where 0 represents a constant number of connections, with no dependence on the amount of computation performed, and 1.0 represents a circuit in which every computation is visible on a connection. Human designed computational systems occupy almost the full range, from spreadsheets in which every computation is visible, to largely serial systems in which minimizing communication (pins) is critical. This relationship is shown in Figure 11. However, when the overriding criterion is that the system must be packed as tightly as possible, Rent observed that the exponent of the power law falls in a close range of roughly 0.5-0.7.

**Figure 11:**
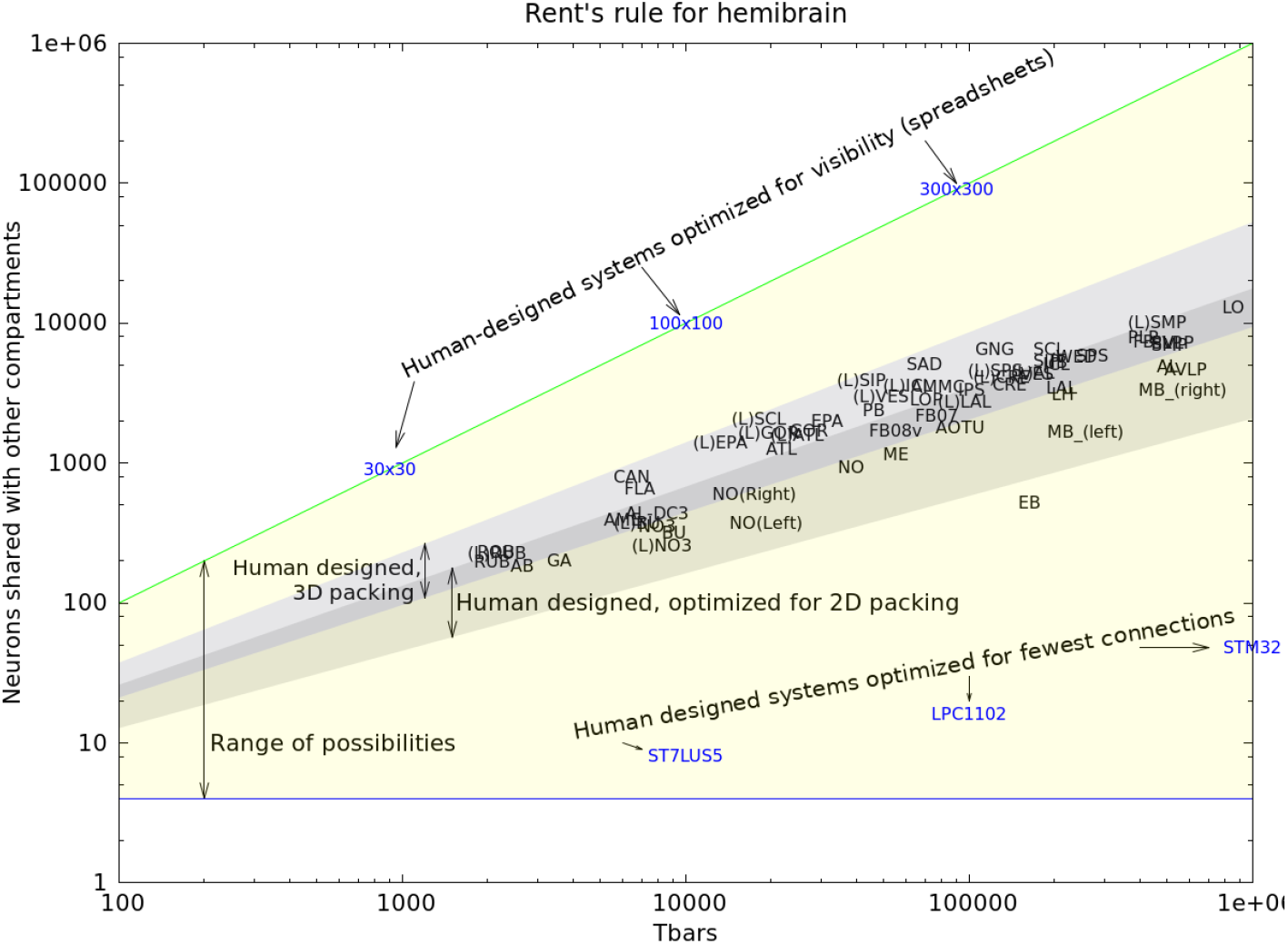
Rent’s rule for the hemi-brain. The yellow region is the theoretical bounds for computation. Human systems designed for visibility into computation achieve the upper bound, while human designed systems designed for minimum communication approach the lower bounds (Microprocessors ST7LU55, LPC1102, and STM32). Human designed systems where efficient packing is the main criterion occupy the shaded area (in 2D and 3D). The hemi-brain compartments fall very nearly in the same range as human designed systems.

For electrical circuits, the computation is measured in gates, and the connections are measured by pin count. These ranges are shown in Figure 11 for circuits that are roughly the size of the fly’s brain, packed in either two[22] or three[23] dimensions.

Also shown in this plot are the values for the fly’s brain computational regions. In this case, the computation is measured as the number of contained T-bars, and the connection count is the number of neurons that have at least one synapse both inside and outside the compartment. (Very similar results are obtained if the computation is measured as the number of PSDs, or the number of unique connection pairs). Almost all the fly brain compartments fall well within the range of exponents expected for packing-dominated systems, while the ellipsoid body (EB) falls just outside the expected area. This is perhaps due to the large number of clique-containing circuits in the ellipsoid body (see Table 3), since such circuits have few connections for the amount of synapses they contain.

Both human designed and biological systems have huge incentives to pack their computation as tightly as possible. A tighter packing of the same computation yields faster operation, lower energy consumption, less material cost, and lower mass. A natural speculation, therefore, is that both the human-designed and evolved systems are dominated by packing considerations, and that both have found similar solutions.

## 3 Conclusions and future work

The main conclusions can be summarized:

- The fly brain neurons have many inputs and outputs (average of 153 of each in our sample).
- The brain is confirmed to be not wired randomly.
- The computational compartments differ significantly in all measured characteristics.
- Path in the fly brain are short, much sorter than in man-made circuits.
- No single input or output dominates.
- The distribution of connection strengths looks like a power law with an exponential tail. This is different from measured connnections in mammalian systems.
- Strong connections are not uniformly distributed.
- Cliques and many-to-one motifs are present in varying numbers.
- The electrical response is strongly determined by the compartment structure.
- There is strong evidence for efficient packing as an important criteria of fly brain evolution.

Further research should involve the whole brain and nervous system, not just the central brain - both brain halves, the ventral nerve cord, and the optic lobes. Additional computational motifs can doubtless be found in the data. Looking at additional specimens of *Drosophila*, and better yet other species, will reveal which finding here are restricted to flies, or perhaps insects, or even invertebrates, and which are more universal. Neuromodulators and their receptors, gap junctions, glia, and unseen cell biology, all are known to affect the function of the brain, but are not included in this connectome. Studies that include these contributions are needed.

## 4 Acknowledgments

We thank our colleagues at Janelia and the broader connectomics field for many helpful discussions and suggestions during the course of this work. Funding for this research was provided by the Howard Hughes Medical Institute.

